# Genome-wide transcriptome analysis reveals the diversity and function of long non-coding RNAs in dinoflagellates

**DOI:** 10.1101/2023.06.27.546665

**Authors:** Yibi Chen, Katherine E. Dougan, Quan Nguyen, Debashish Bhattacharya, Cheong Xin Chan

## Abstract

Dinoflagellates are a diverse group of phytoplankton, ranging from harmful bloom-forming microalgae to photosymbionts that are critical for sustaining coral reefs. Genome and transcriptome data from dinoflagellates are revealing extensive genomic divergence and lineage-specific innovation of gene functions. However, most studies thus far have focused on protein-coding genes; long non-coding RNAs (lncRNAs), known to regulate gene expression in eukaryotes, are largely unexplored. Here, using both genome and transcriptome data, we identified a combined total of 48,039 polyadenylated lncRNAs in the genomes of three dinoflagellate species: the coral symbionts of *Cladocopium proliferum* and *Durusdinium trenchii*, and the bloom-forming *Prorocentrum cordatum*. These putative lncRNAs are shorter, and have fewer introns and lower G+C-content when compared to protein-coding sequences. Although 37,768 (78.6%) lncRNAs shared no significant similarity with one another, we classified all lncRNAs based on conserved sequence motifs (*k*-mers) into distinct clusters following properties of potential protein-binding and/or subcellular localisation. Interestingly, 3708 (7.7%) lncRNAs were differentially expressed in response to heat stress, lifestyle, and/or growth phases, and they shared co-expression patterns with protein-coding genes. Based on inferred triplex interactions between lncRNA and upstream (putative promoter) regions of protein-coding genes, we identified a combined 19,460 putative gene targets for 3,721 lncRNAs; 907 genes exhibit differential expression under heat stress. These results reveal for the first time the functional diversity of lncRNAs in dinoflagellates, and demonstrate how lncRNAs, often overlooked in transcriptome data, could regulate gene expression as a molecular response to heat stress in these ecologically important organisms.

## Introduction

Dinoflagellates are a diverse group of microbial eukaryotes found ubiquitously in both marine and freshwater environments. These taxa range from parasites, symbionts of various organisms in coral reefs, to free-living species, of which some form harmful algal blooms that have serious implications for human health. Dinoflagellates are known to have large, complex genomes up to ∼200 Gbp in size, with features atypical of eukaryotes and packaged in permanently condensed chromosomes (Hou and Lin 2009; Lin 2011; Wisecaver and Hackett 2011). The available genome data from dinoflagellates have revealed extensive genomic divergence and the contribution of lifestyle to functional and structural diversification of genes and genomic elements (John et al. 2019; González-Pech et al. 2021; Dougan et al. 2022b), including distinct phylogenetic signals between coding and non-coding sequence regions (Lo et al. 2022).

Earlier studies of transcriptome data identified: (a) differentially expressed genes in dinoflagellates relative to lifestyle (Bellantuono et al. 2019), and to biotic and abiotic stresses (Moustafa et al. 2010; Lowe et al. 2011; Wang et al. 2019; Kang et al. 2021), (b) the *trans*-splicing of spliced-leader sequences at the 5′-end of mature mRNAs (Zhang et al. 2007), (c) diversity of transcript isoforms (Chen et al. 2022a), and, more recently in combination with genome data, (d) protein-coding sequences (Shoguchi et al. 2013; Lin et al. 2015; Aranda et al. 2016; Shoguchi et al. 2018; Chen et al. 2020; Stephens et al. 2020; González-Pech et al. 2021; Chen et al. 2022b), and (e) the differential editing of mRNAs (Liew et al. 2017; Dougan et al. 2022b) and differential exon usage (Dougan et al. 2022a) in distinct growth conditions. In most cases, these studies employed standard RNA-Seq approaches for eukaryotes to generate transcriptome data, whereby mRNAs with polyadenylated tails are retained, and focused on protein-coding genes. Many non-coding RNAs (ncRNAs) would have retained polyadenylated tails following transcription similarly to mRNAs (Ip and Nakagawa 2012); they are likely also present in these transcriptome datasets but were largely ignored.

Our current understanding of how gene expression is regulated in dinoflagellates remains limited. In eukaryotes, transcription factors (TFs) bind to specific DNA region in the genome, which could enhance or supress the transcription of associated gene(s). TFs make up ∼4% of the proteome in yeast, and ∼8% of the proteome in plants and mammals (Babu et al. 2004). In comparison, proteins that contain DNA-binding domains were found to account for only 0.15%–0.30% of all proteins in dinoflagellates (Bayer et al. 2012; Beauchemin et al. 2012). In an independent analysis of the dinoflagellates *Lingulodinium polyedra* and *Fugacium kawagutii*, ∼60% of putative TFs (i.e. DNA-binding proteins) contain a cold shock domain, and they tend to bind to RNA instead of specific double-stranded DNA as expected for canonical TFs (Zaheri and Morse 2021). This observation hints at the role of RNAs in regulating gene expression in dinoflagellates.

The regulatory role of ncRNAs in gene expression has been demonstrated in diverse eukaryotes (Hombach and Kretz 2016; Mattick et al. 2023). The two most common types of ncRNAs with regulatory functions are microRNAs (21-23 nt in length) and long non-coding RNAs (lncRNAs; >200 nt (Bartel 2009; Kopp and Mendell 2018), more recently recommended as >500 nt (Mattick et al. 2023)). Target mRNA molecules are bound by microRNAs, which supress translation of the transcript into protein and/or cause their degradation. In dinoflagellates, microRNAs have been associated with the regulation of protein modification, signalling, gene expression, sulfatide catabolism (Shi et al. 2017), and amino acid metabolism (Yu et al. 2020), as well as the molecular responses to DNA damage (Baumgarten et al. 2013) and symbiosis (Lin et al. 2015). In contrast, lncRNAs in eukaryotes are known to serve as molecular scaffolds that link TFs to target genes, thereby altering their transcription (Moison et al. 2021). The genome loci for lncRNA binding have also been associated with the recruitment of chromatin-modifying complexes, thus changing the accessibility of nearby genes or the entire chromosomes (Cerase et al. 2015). To date, lncRNAs in dinoflagellates remain uncharacterised and largely unexplored.

Here, we investigate the conservation and functional role of lncRNAs in dinoflagellates relative to distinct growth conditions using high-quality genome assemblies and transcriptome datasets from three taxa: *Cladocopium proliferum* SCF055 (Camp et al. 2022; Chen et al. 2022b) (an isolate formerly identified as *Cladocopium goreaui* SCF055 (Butler et al. 2023)), *Durusdinium trenchii* CCMP2556 (Bellantuono et al. 2019; Dougan et al. 2022a), and *Prorocentrum cordatum* CCMP1329 (Dougan et al. 2022b), all generated under heat-stress experiments (Table 1). Due to the different experimental designs associated with these independently generated datasets, they allow for assessment of heat-stress response at slightly different resolutions: (a) early-versus late-stage response (*C. proliferum*), (b) free-living versus symbiotic stages (*D. trenchii*), and (c) exponential versus stationary growth phases (*P. cordatum*).

**Table 1.**
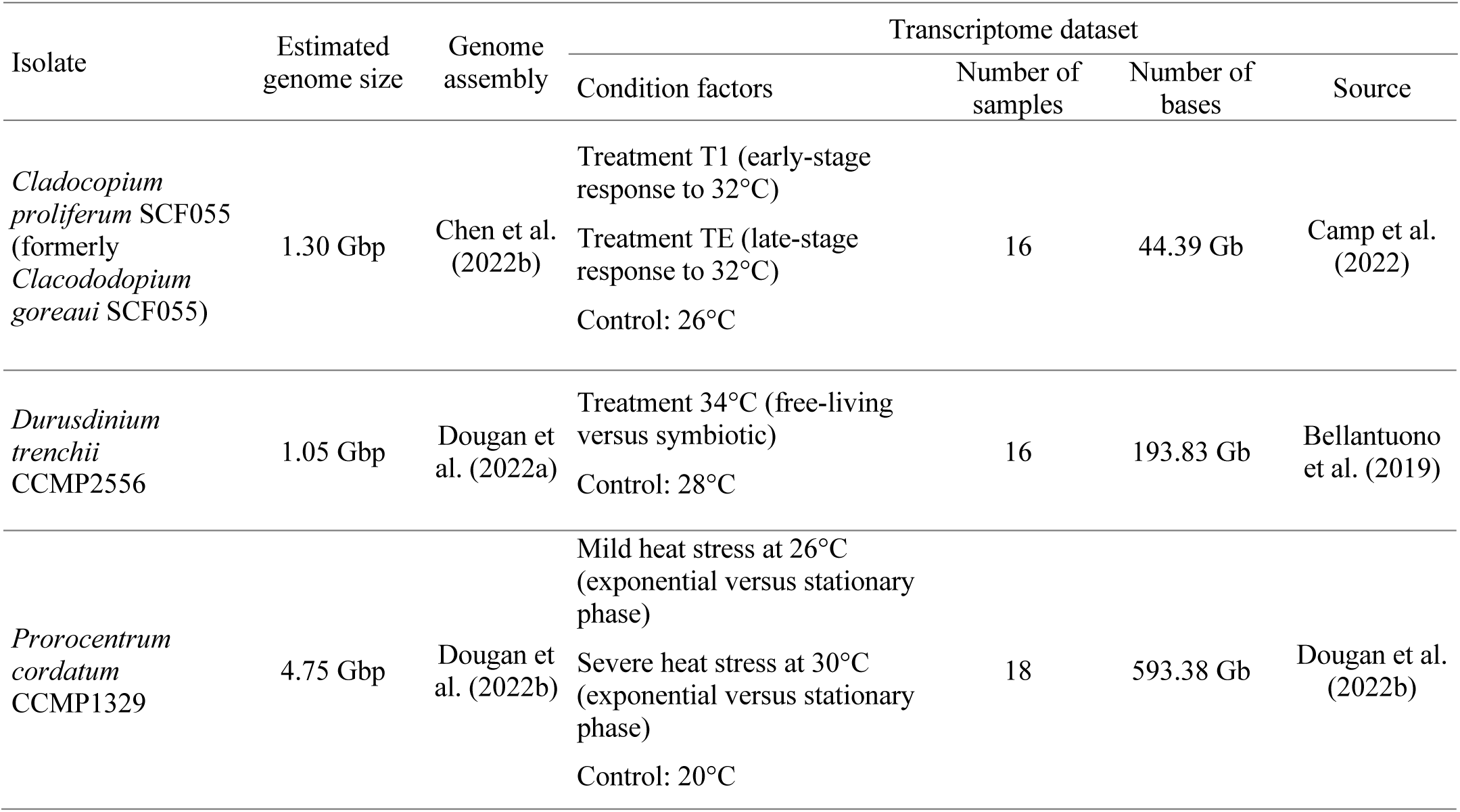
The three core dinoflagellate transcriptome datasets used in this study.

## Results and Discussion

### LncRNAs have fewer introns and lower G+C-content than protein-coding genes

We adopted a stringent approach to identify putative lncRNAs, focusing on expressed RNAs transcribed from genome regions that do not overlap with exons. For each of the three core transcriptome datasets (Table 1), we first assembled the transcriptome independently, from which we identified the lncRNAs by incorporating structural annotation of the reference genome sequences from the corresponding taxa. We consider a transcript to be lncRNA if it (a) mapped to the reference genome (at ≥95% sequence identity covering ≥75% of query) and (b) did not encode any Pfam protein domains, was (c) classified as non-coding by three methods for calculating protein-coding potential, and (d) expressed in ten or more samples; see Methods for detail. For clarity of presentation, hereinafter we define *gene* strictly as a protein-coding gene, excluding lncRNAs.

Using this approach, we identified distinct lncRNAs in *C. proliferum* (13,435), *D. trenchii* (7,036), and *P. cordatum* (27,568), with a combined total of 48,039 (Table 2). The number of lncRNAs in *P. cordatum* is more than twice of that in *C. proliferum* (genome size ∼1.5 Gbp, 45,322 genes) (Chen et al. 2022b), likely due to the larger genome size (∼4.75 Gbp) and number of genes (85,849) in *P. cordatum* (Dougan et al. 2022b). However, proportionately, the number of lncRNAs is approximately one-third of that of genes in these two genomes, i.e. 0.30 in *C. goreaui* and 0.32 in *P. cordatum* (Table 2). The equivalent ratio is much smaller (0.13) in *D. trenchii*, which may be affected by whole-genome duplication implicated in this lineage (Dougan et al. 2022a).

**Table 2.** The lncRNAs identified in the three dinoflagellate taxa, their statistics relative to genes and the associated lncRNA:gene ratios.

| Taxon | *Cladocopium proliferum*<br>SCF055 |  |  | *Durusdinium trenchii*<br>CCMP2556 |  |  | *Prorocentrum cordatum*<br>CCMP1329 |  |  |
| --- | --- | --- | --- | --- | --- | --- | --- | --- | --- |
| Metric | lncRNA | Gene | Ratio | lncRNA | Gene | Ratio | lncRNA | Gene | Ratio |
| Counts | 13,435 | 45,322 | 0.30 | 7,036 | 55,799 | 0.13 | 27,568 | 85,849 | 0.32 |
| Mean length (bp) | 450 | 2,018 | 0.22 | 447 | 1,647 | 0.27 | 493 | 2,798 | 0.18 |
| % G+C | 48.8 | 54.2 | 0.90 | 51.7 | 55.7 | 0.93 | 61.7 | 65.9 | 0.94 |
| Mean number of introns per lncRNA/gene | 0.8 | 16.3 | 0.05 | 2.0 | 15.7 | 0.13 | 0.39 | 9.8 | 0.04 |
| % lncRNAs/genes that contain introns | 23.0 | 95.9 | 0.24 | 51.7 | 93.1 | 0.56 | 27.3 | 83.7 | 0.33 |

Among the three genomes, lncRNAs (mean length 447–493 bp) were shorter than the genes (mean length 1,657–2,798 bp; Table 2). These lncRNA sequences are shorter than commonly expected (∼1 to >100 Kbp) (Mattick et al. 2023), likely due to the RNA-Seq (short-read) transcriptome datasets used in this study, in which full-length lncRNAs may not be recovered as readily as the long-read technology (Palos et al. 2022), and non-polyadenylated lncRNAs (Mattick et al. 2023) would have been excluded. The lncRNAs exhibited lower (0.90–0.94 fold) G+C-content that did the genes (e.g. mean 48.8% compared to 54.2% for *C. proliferum*; Table 2), and had fewer introns, e.g. mean 0.39 introns per lncRNA compared to mean 9.8 introns per gene of *P. cordatum* (Table 2). These numbers are comparable to earlier studies of eukaryotes, e.g. lncRNAs among the Brassicaceae plants are shorter and with lower G+C-content compared to the genes (Palos et al. 2022), and those in zebra finch have fewer introns than do the genes (Chen et al. 2017).

### LncRNAs exhibit conserved motifs despite sharing low overall sequence similarity

To assess conservation of lncRNAs, we first identified 2,325 putatively homologous sets from the 48,039 lncRNAs recovered in all three taxa based on shared sequence similarity. Most lncRNAs (37,768; 78.6%) were not assigned to any homologous sets (i.e. they are singletons). Among those assigned to a set, most (2291 of 2325 [98.5%] sets) represent highly duplicated lncRNAs within a genome, e.g. 1,559 homologous sets implicating 8,063 lncRNAs are specific to *P. cordatum* (Figure 1a). This result is consistent with the expectation that, in contrast to genes, lncRNA sequences are not highly conserved across taxa (Johnsson et al. 2014). Few exceptions are however known, such as the MALAT1 lncRNA with a highly conserved 3′-end processing module in animals and a functional role in cell cycle progression and cell migration in humans (Diederichs 2014). Interestingly, two homologous sets implicating 24 lncRNAs were identified in all three taxa. Given the lack of lncRNA resources for non-model systems, we searched these lncRNAs against the high-quality curated database of human lncRNAs, FANTOM CAT (Hon et al. 2017) and other lncRNAs from the NCBI nr database. One of the two sets, containing 19 lncRNAs, show significant sequence similarity (BLASTn, *E* ≤ 10^-5^) with the human lncRNA CATG00000009539.1 in FANTOM CAT, and the *Mus musculus* lncRNA MSUR1 in the nr database; CATG00000009539.1 is an intergenic lncRNA expressed in small intestine cells (Lizio et al. 2015), whereas MSUR1 is known to rescue cell death mediated by copper/zinc superoxide dismutase (Kaur et al. 2013). The other set does not share significant sequence similarity to any sequences in the databases. Although the function of these lncRNAs in dinoflagellates remains to be validated experimentally, the observed sequence similarity (e.g. 79.78% identity shared between lncRNA of *P. cordatum* over 75.43% of the MSUR1 sequence) suggests some extent of cross-phylum sequence conservation.

**Figure 1.**
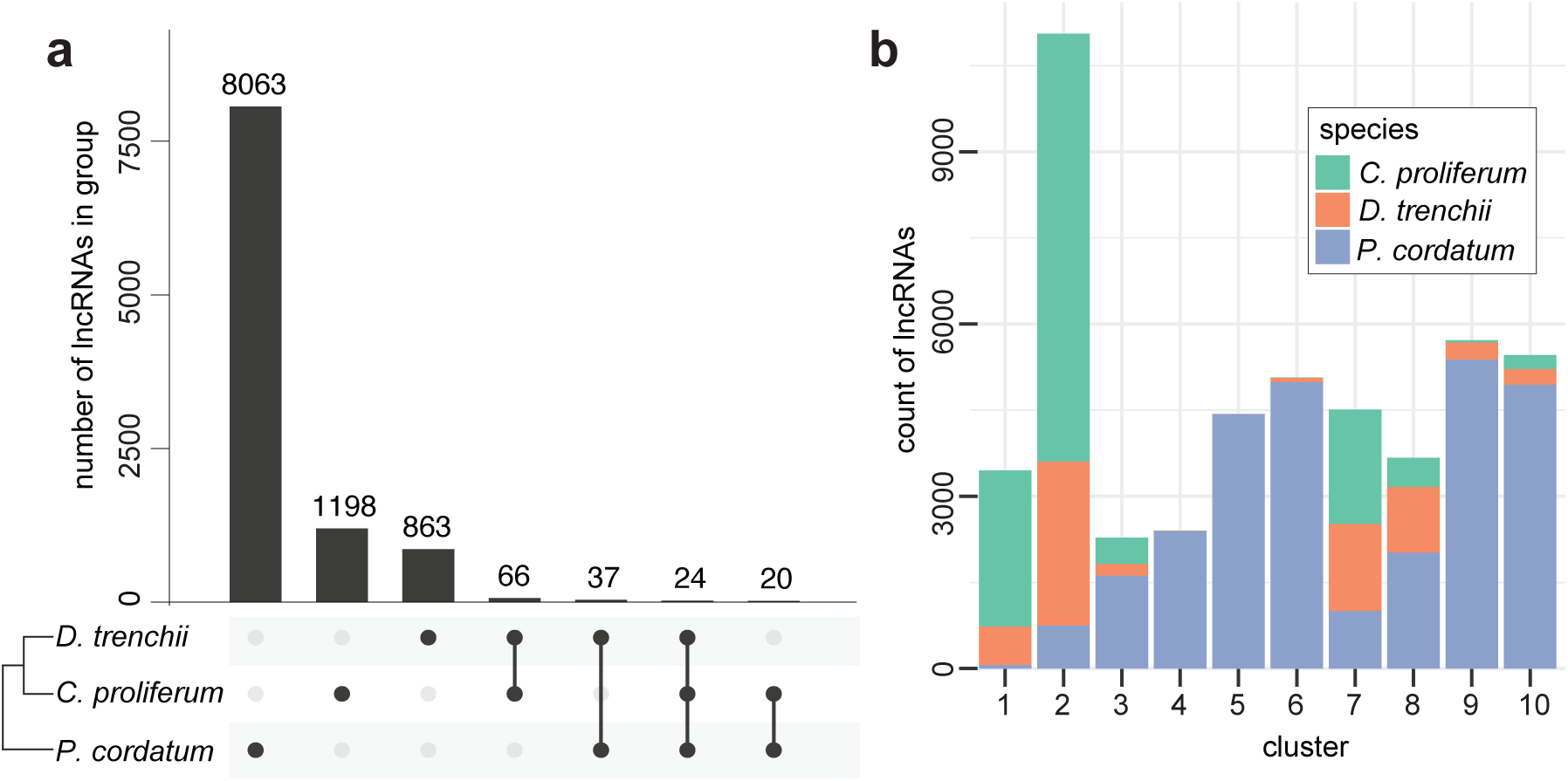
Homologous lncRNAs identified in the three dinoflagellate taxa based on (a) shared sequence similarity, and (b) conserved sequence motifs of *k*-mers.

Our results lend support to the lack of shared sequence similarity and/or contiguity among lncRNAs, as observed among homologous lncRNAs in other eukaryotes (Li and Yang 2017). For this reason, the alignment-free approach based on conserved *k*-mers (i.e. short motifs of length *k*) provides a good alternative for identifying lncRNAs with similar functions, specifically those containing motifs specific to protein binding and subcellular localisation (Kirk et al. 2018). Using this approach, we generated normalised *k*-mer profiles using all lncRNAs from the three genomes as background and calculated a pairwise adjacency matrix for all possible lncRNA pairs. Using hierarchical clustering, we classified all lncRNAs into ten clusters (Figure 1b; see Methods); those from *C. proliferum* and *D. trenchii* tend to group together (e.g. clusters 1 and 2), distinct from *P. cordatum* (e.g. clusters 4, 5, and 6). This observation suggests more conserved lncRNA motifs for putative protein binding and subcellular localisation between the coral symbionts *C. proliferum* and *D. trenchii* (both of Family Symbiodiniaceae in Order Suessiales), compared to the free-living *P. cordatum* (Order Prorocentrales), consistent with phylogenetic relationships among the three taxa.

### LncRNAs are differentially expressed in response to heat stress

Using the transcriptome dataset for each taxon, we identified differentially expressed (DE) lncRNAs in response to heat stress, growth phase, and/or lifestyle (Figure 2), based on a stringent criterion (see Methods); in combination, we observed 3,708 DE lncRNAs. In *C. proliferum*, no DE lncRNAs were observed during early-stage heat stress (T1) relative to controls (Figure 2a) compared to 139 DE lncRNAs at late-stage heat stress (TE; Figure 2b), a pattern which is consistent with the DE genes (1 in Figure 2a, 146 in Figure 2b). Similarly, in an experiment that measured heat-stress response of *D. trenchii* at day 3 of temperature ramping to 34°C, few DE lncRNAs (7) and genes (11) were observed (combining free-living and symbiotic stages; Figure 2c). Interestingly, we identified three orders of magnitude greater DE lncRNAs (1,572) and genes (3,669) under heat stress in *P. cordatum* (combining both exponential and stationary growth phases; Figure 2d) than those observed for the two symbiotic species. However, whether this stark difference was related to the distinct ecological niches among the three taxa is unclear due to the different experimental designs from which the data were generated, including their sequencing depths. We also observed DE lncRNAs relative to free-living versus symbiotic stages in *D. trenchii* (Figure 2e), and to distinct growth phases of *P. cordatum* (Figure 2f), as similarly observed for the DE genes in the earlier studies (Bellantuono et al. 2019; Dougan et al. 2022a). A greater transcriptional response to heat stress was observed in *P. cordatum* cells during stationary growth phase (longer exposure to heat stress) than those in exponential phase (shorter exposure to heat stress). Our results, in combination with the previous observations (Camp et al. 2022; Dougan et al. 2022b), suggest a greater transcriptional response in dinoflagellates under prolonged heat stress. This may explain in part why in some earlier studies (Barshis et al. 2014; Roy et al. 2014) that assessed transcriptional changes within a short period found few DE genes in dinoflagellates.

**Figure 2.**
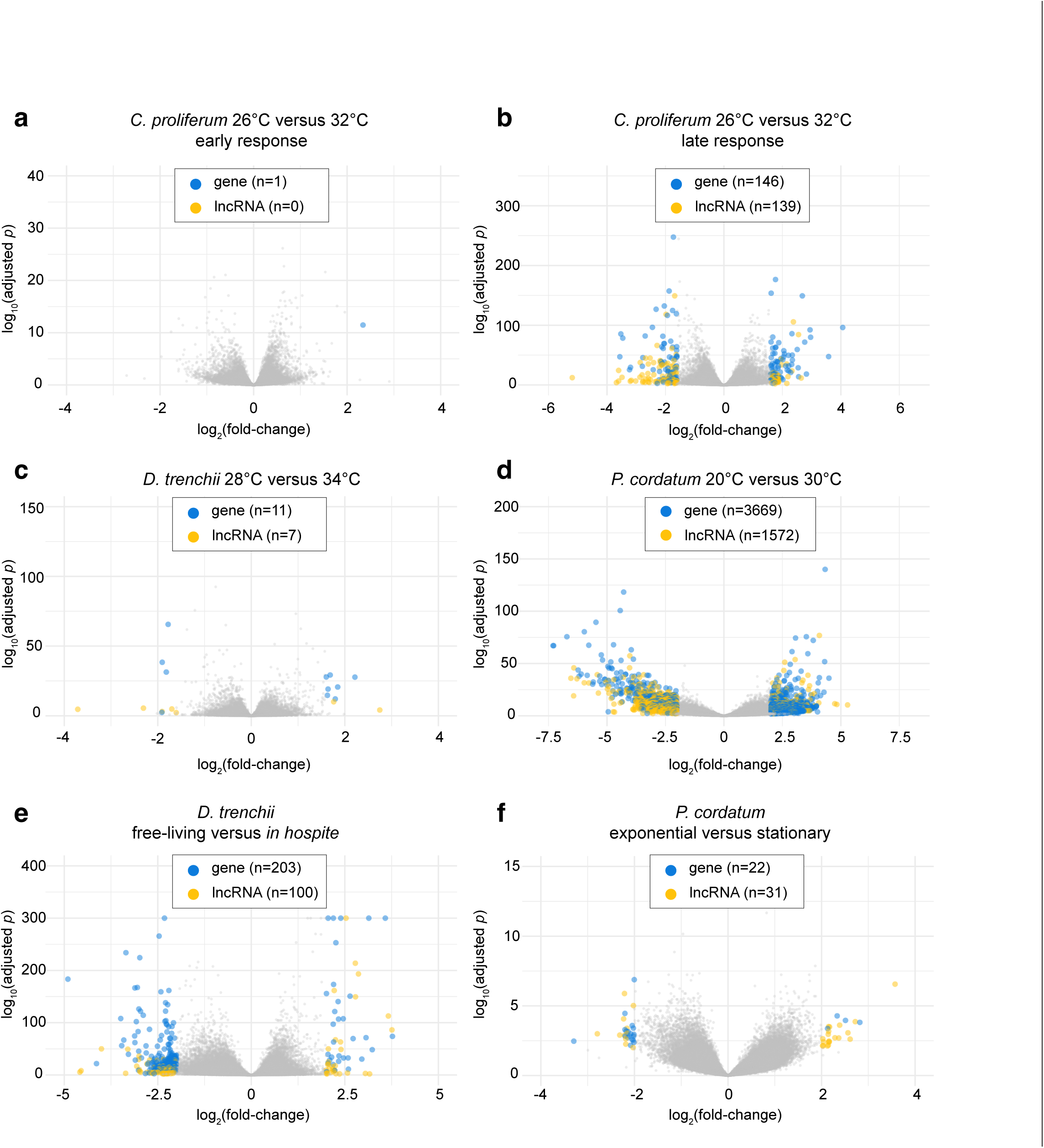
Differentially expressed genes and lncRNAs relative to heat stress, lifestyle, and growth phase. Volcano plots are shown for (a) *C. proliferum* (early-stage response T1: 32 versus 26°C), (b) *C. proliferum* (late-stage response TE: 32 versus 26°C), (c) *D. trenchii* (34 versus 28°C), (d) *P. cordatum* (30 versus 20°C), (e) *D. trenchii* (free-living versus symbiotic stage), and (f) *P. cordatum* (exponential versus stationary phase). For each panel, the *x*-axis represents fold-change of transcript expression (in log_2_ scale), and the *y*-axis represents the significance of difference in adjusted *p*-value (in log_10_ scale). Differentially expressed genes (blue) and differentially expressed lncRNAs (yellow) were noted, and their numbers are shown for each panel.

### LncRNAs of similar *k*-mer profiles tend to be co-expressed

We used weighted gene co-expression network analysis (WGCNA) to assess co-expression of genes and lncRNAs in the three transcriptome datasets (see Methods). A WGCNA module represents a group of similarly co-expressed genes and lncRNAs, implicating biological processes or metabolic functions that share a similar molecular response. Independently, we identified WGCNA modules for *C. proliferum* (41; Figure 3a), *D. trenchii* (11; Figure 3b) and *P. cordatum* (53; Figure 3c). The number of lncRNAs within each module shows a strong correlation with the number of genes (Spearman correlation coefficient = 0.88, 0.97, and 0.81 for *C. proliferum*, *D. trenchii*, and *P. cordatum*). This observation suggests that WGCNA modules containing a greater number of genes, which likely participate in more complex biological processes and/or metabolic pathways, tend to contain a larger number of lncRNAs.

**Figure 3.**
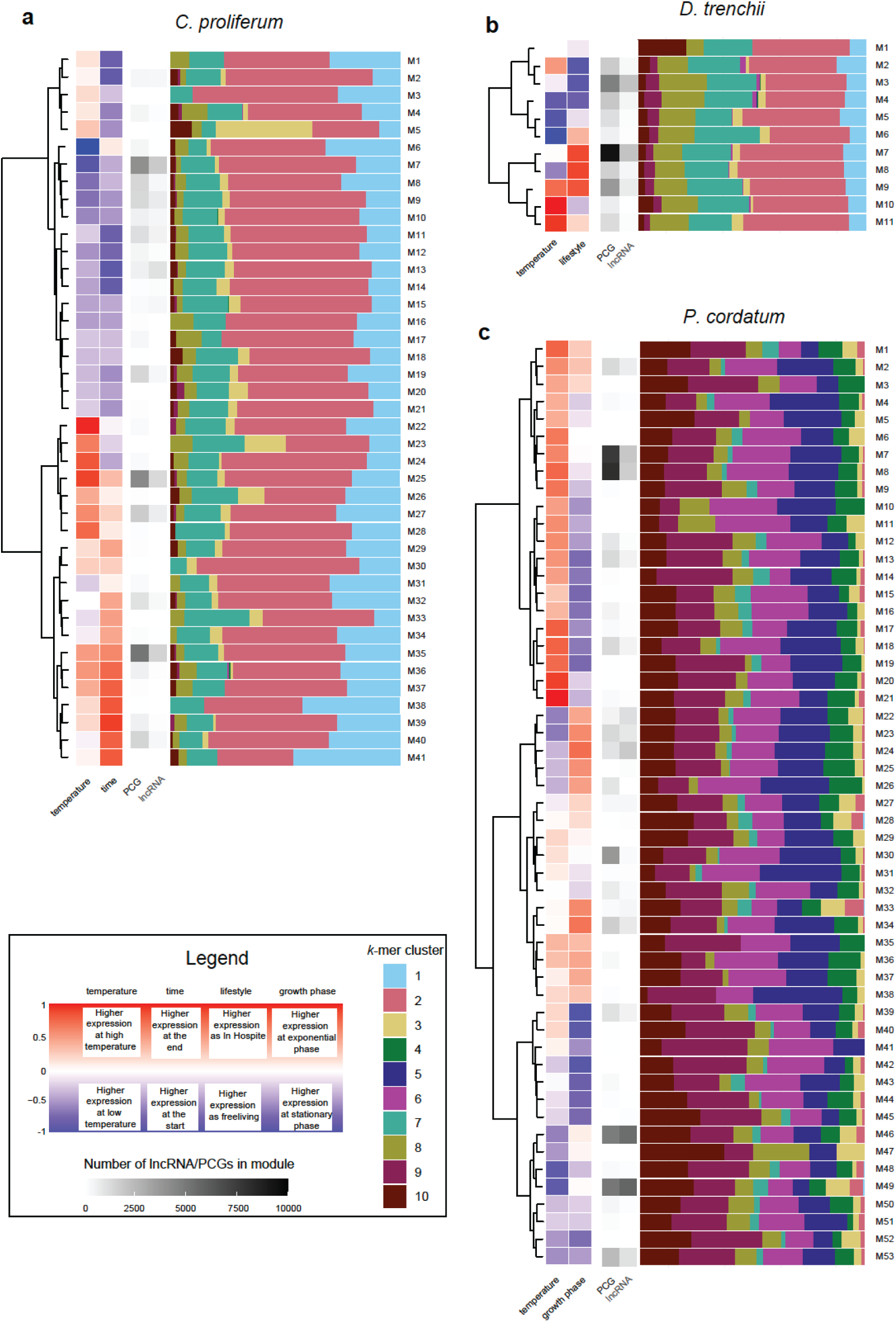
WGCNA modules of co-expressed lncRNAs, shown for (a) *C. proliferum*, (b) *D. trenchii*, and (c) *P. cordatum*. In each panel, each row represents a WGCNA module, showing, from left to right, the Pearson correlation coefficient of eigengenes to the two external factors examined in the dataset (temperature, and time/lifestyle/growth phase), the number of genes and lncRNAs in the module, and the proportion of all lncRNAs based on *k*-mer cluster in stacked bar charts.

The *k*-mer profiles of lncRNAs have been associated with the repression or activation of nearby genes in eukaryotes, and with lncRNAs sharing similar protein-binding properties, linking conserved motifs to lncRNA function (Kirk et al. 2018). In each dinoflagellate genome, we observed a significant correlation (Fisher’s exact test, 10,000-replicate Monte Carlo simulations, *p* <10^-5^) between the overall distribution of *k*-mer clusters and the expression pattern based on WGCNA modules. Among the WGCNA modules for each taxon, we identified significant differential representation of one or more *k*-mer clusters in *C. proliferum* (12 of 41; Table <u>S1</u>), *D. trenchii* (2 of 11; Table <u>S2</u>), and *P. cordatum* (17 of 53; Table <u>S3</u>). For example, in *P. cordatum*, *k*-mer clusters 5 and 6 were overrepresented in the two largest modules up-regulated under heat stress (M7 and M8) but underrepresented in the two largest modules down-regulated under heat stress (M46 and M49); the opposite trend was observed for *k*-mer clusters 2, 3, and 10 (Figure 3c and Table <u>S3</u>). The conserved motifs of these implicated lncRNAs are rich in pyrimidine nucleotides (cytosine and thymidine; Figure S1), indicating their capacity of binding to purine-rich DNA sequences via Hoogsteen (or reverse Hoogsteen) hydrogen bonds (Li et al. 2016). This mode of action is expected in the mechanism that underpins lncRNA regulation of gene expression in eukaryotes, whereby lncRNAs form a triple-helical structure by binding to the purine-rich strand of target genome regions, e.g. promoters (Sentürk Cetin et al. 2019).

These results clearly indicate a correlation between conserved lncRNA motifs and gene expression, and by extension, gene function in dinoflagellates. Although the specific binding sites and regulatory functions of these lncRNAs remain to be experimentally validated, considering the binding affinity of putative TFs to RNA molecules in dinoflagellates (Zaheri and Morse 2021), these results demonstrate the potential role of ncRNAs in regulating gene expression in these species.

### Gene targets of lncRNAs based on the predicted formation of triple-helical structures

We identified putative gene targets for all lncRNAs by inferring triplex interactions between each lncRNA and the upstream (promoter) region of a gene, following Buske et al. (2012); see Methods. We identified lncRNAs in *C. proliferum* (168; 1.3% of 13,435), *D. trenchii* (142; 2.0% of 7,036), and *P. cordatum* (3,411; 1.2% of 27568) that putatively form a triple-helix with promoter regions of 439, 622 and 18,399 genes (Table 3). Based on the specificity of interactions between lncRNAs to genes, we categorised these lncRNAs broadly into four groups: one-to-one, one-to-many, many-to-one, and many-to-many. Among these groups, no lncRNAs showed many-to-one interactions with genes in all cases. Many-to-many interactions were the most abundant, comprising 58.9%, 60.6%, and 82.7% of triplex-forming lncRNAs in *C. proliferum*, *D. trenchii*, and *P. cordatum*, followed by the group of one-to-one interactions, targeting genes encoding distinct functions of biological processes (Tables <u>S4</u> through <u>S9</u>). For instance, relative to all genes of *C. proliferum*, those implicated in many-to-many interactions (Table <u>S4</u>) appear to be enriched in functions related to photoreactive repair and vesicle-mediated transport based on annotation of Gene Ontology (GO) terms, whereas those in one-to-one interactions (Table <u>S5</u>) are enriched in functions related to methylation and post-replication repair.

**Table 3.** Predicted triplex interactions of lncRNAs and protein-coding genes in the three dinoflagellate datasets. DE genes in response to heat stress were shown for *C. proliferum* (26 versus 32°C), *D. trenchii* (28 versus 34°C), and *P. cordatum* (20 versus 30°C).

|  | *C. proliferum* | *D. trenchii* | *P. cordatum* |
| --- | --- | --- | --- |
| Number of triplex-forming lncRNAs | 168 | 142 | 3411 |
| Number of interactions | 1085 | 2414 | 288977 |
| One-to-one | 59 | 39 | 558 |
| One-to-many | 10 | 17 | 33 |
| Many-to-one | 0 | 0 | 0 |
| Many-to-many | 99 | 86 | 2820 |
| Total number of genes | 45,322 | 55,799 | 85849 |
| Genes with interacting lncRNAs | 439 (0.9%) | 622 (1.1%) | 18399 (21.4%) |
| Total number of DE genes under heat stress | 146 | 11 | 3669 |
| DE genes under heat stress with interacting lncRNAs | 3 (2.1%) | 0 | 904 (24.6%) |

We found a greater proportion of genes interacting with lncRNAs in the free-living *P. cordatum* (21.4%), compared to the two coral symbionts *C. proliferum* (0.9%) and *D. trenchii* (1.1%) (Table 3). Moreover, this proportion is even greater for DE genes under heat stress (24.6%), and the same trend is also observed in *C. proliferum* (2.1% DE genes versus 0.9% all genes; Table 3). Interestingly, we observed no lncRNA interactions implicating DE genes associated with heat-stress response in *D. trenchii* (Table 3); this may be due in part to the fact that *D. trenchii* is a naturally thermotolerant species for which the distinct lifestyle (free-living versus symbiotic) stages elicited a much stronger differential molecular response than did heat stress (Bellantuono et al. 2019; Dougan et al. 2022a). Although we cannot dismiss potential false positives in our prediction of triplex interactions (Antonov et al. 2019), particularly among the one-to-one interactions, our results indicate lncRNA interactions with target genes as a molecular response to heat stress in dinoflagellates.

To assess the impact of lncRNAs on the overall transcriptional response to heat stress, we focused on *P. cordatum* for which the largest number of lncRNAs were identified. Specifically, we investigated the inferred interactions between lncRNAs and DE genes, i.e. lncRNAs for which expression is positively or negatively correlated to the DE genes based on the Spearman correlation coefficient ≥0.8 and *p* ≤ 0.05 following Fan et al. (2019). We identified 2941 interactions of 435 lncRNAs implicating 527 genes; remarkably, of these interactions, 2398 (81.5%) showed a negative correlation with the expression of DE genes, suggesting a suppression of the interacting lncRNAs on expression of target genes. This trend aligns with the current knowledge of triplex-forming ncRNAs, most of which repress gene expression (Martianov et al. 2007; Grote and Herrmann 2013; Grote et al. 2013; Li et al. 2016; Leisegang et al. 2022).

Among the 435 lncRNAs, 186 (42.8%) have only one gene target, implicating 137 genes, whereas the remaining 249 have multiple gene targets, forming 2755 (93.7% of 2941) interactions with 488 genes. The correlation of expression between these 488 genes and their interacting lncRNAs revealed three distinct clusters (Figure S2): Group I (359 genes), Group II (64), and Group III (65). Group I had the least number of lncRNA interactions (mean 2.37), compared to Group II (11.08) and Group III (18.37). To reduce the bias of potential false positives in our interpretation of these results, we focused on Groups II and III for which a greater extent of lncRNA interactions were observed (Figure 4). Interestingly, these groups consist of only up-regulated genes under heat stress. Figure 4 shows a clear pattern of negative correlation between the expression of interacting lncRNAs and the expression of up-regulated genes, and the association of lncRNA interactions to distinct gene functions of Groups II and III. Based on annotation of GO terms, the up-regulated genes in Group III largely encode functions related to microtubule-based movement and polysaccharide metabolic process, compared to those largely encoding functions related to protein metabolic processes and macromolecule modification in Groups I and II. For instance, three genes in Group III encode for the dynein heavy chain proteins (Figure 4) that are main structural and functional components of dynein motor complexes essential for intracellular transport and cell movements in eukaryotes, often encoded in multiple genes (Asai and Wilkes 2004). Up-regulation of these genes has been associated with phosphorus deficiency in *Prorocentrum shikokuense*, indicating enhanced intracellular trafficking or cell motility due to stress (Shi et al. 2017). Our result hints at the role of lncRNAs in regulating the expression of these genes in *P. cordatum* as a heat-stress response.

**Figure 4.**
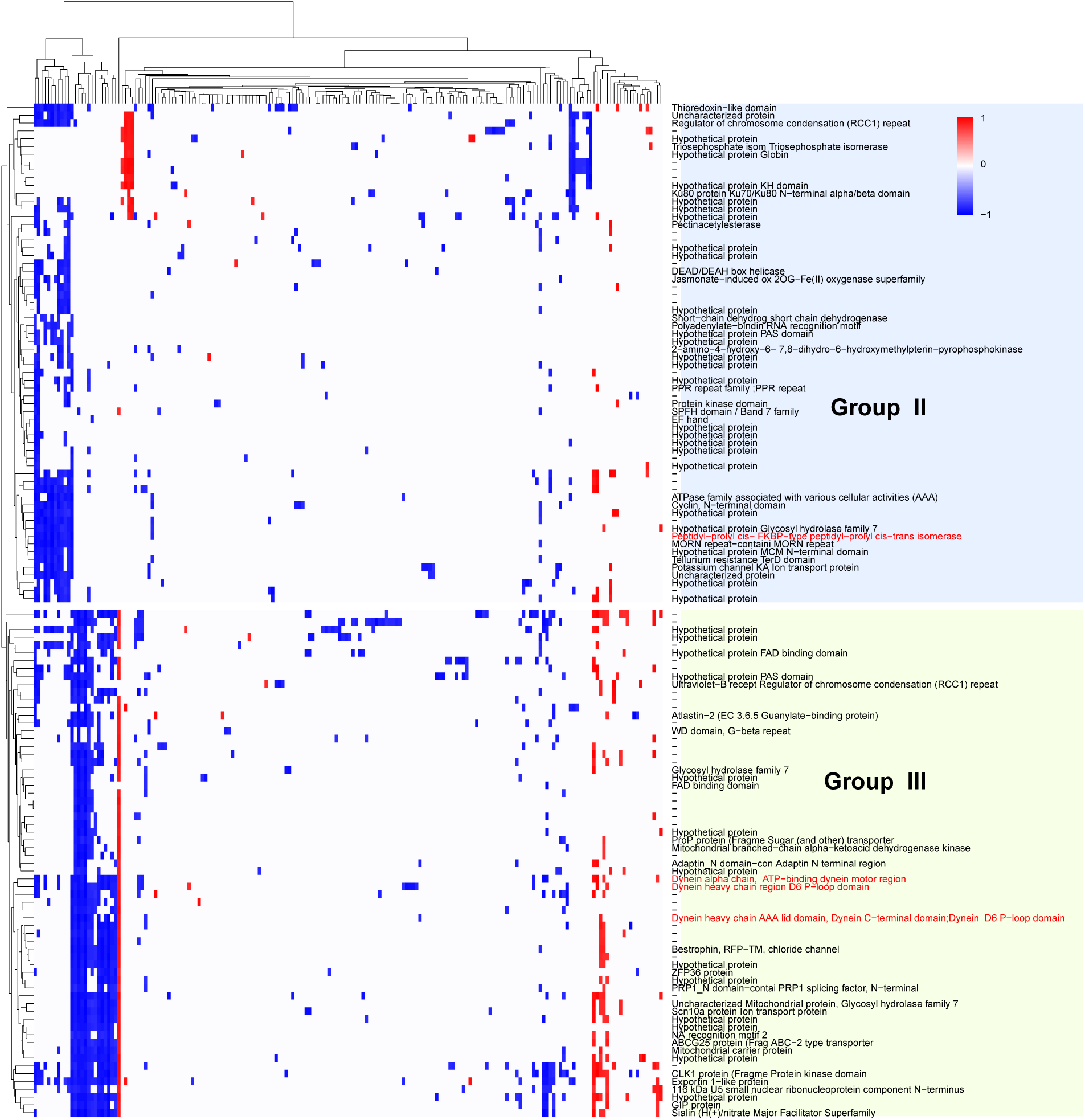
Correlation of expression of up-regulated *P. cordatum* genes under heat stress (30 versus 20°C) with the expression of their interacting lncRNAs. Each row represents a gene, and each column represents a lncRNA. Spearman correlation of expression (absolute value ≥0.8) is shown for each gene-lncRNA pair, with –1.0 indicating negative correlation, and 1.0 indicating positive correlation.

### Hypothesised regulation of transcription initiation by lncRNAs in a heat-stress response

An up-regulated gene in Group II encodes peptidyl-prolyl *cis*-*trans* isomerase (PPIase). PPIase catalyses the interconversion of the *cis* and *trans* isomers of peptidyl-prolyl peptide bonds, which is essential for proper folding of proteins that affects their stability and function (Vallon 2005). This up-regulated gene in *P. cordatum* is the putative target of 13 lncRNAs – that are down-regulated under heat stress (Figure 4). The predicted purine-rich binding site for these lncRNAs is located between 193 bp and 226 bp upstream of the protein-coding sequence (Figure 5a), hypothesised to form the triple helical structure.

**Figure 5.**
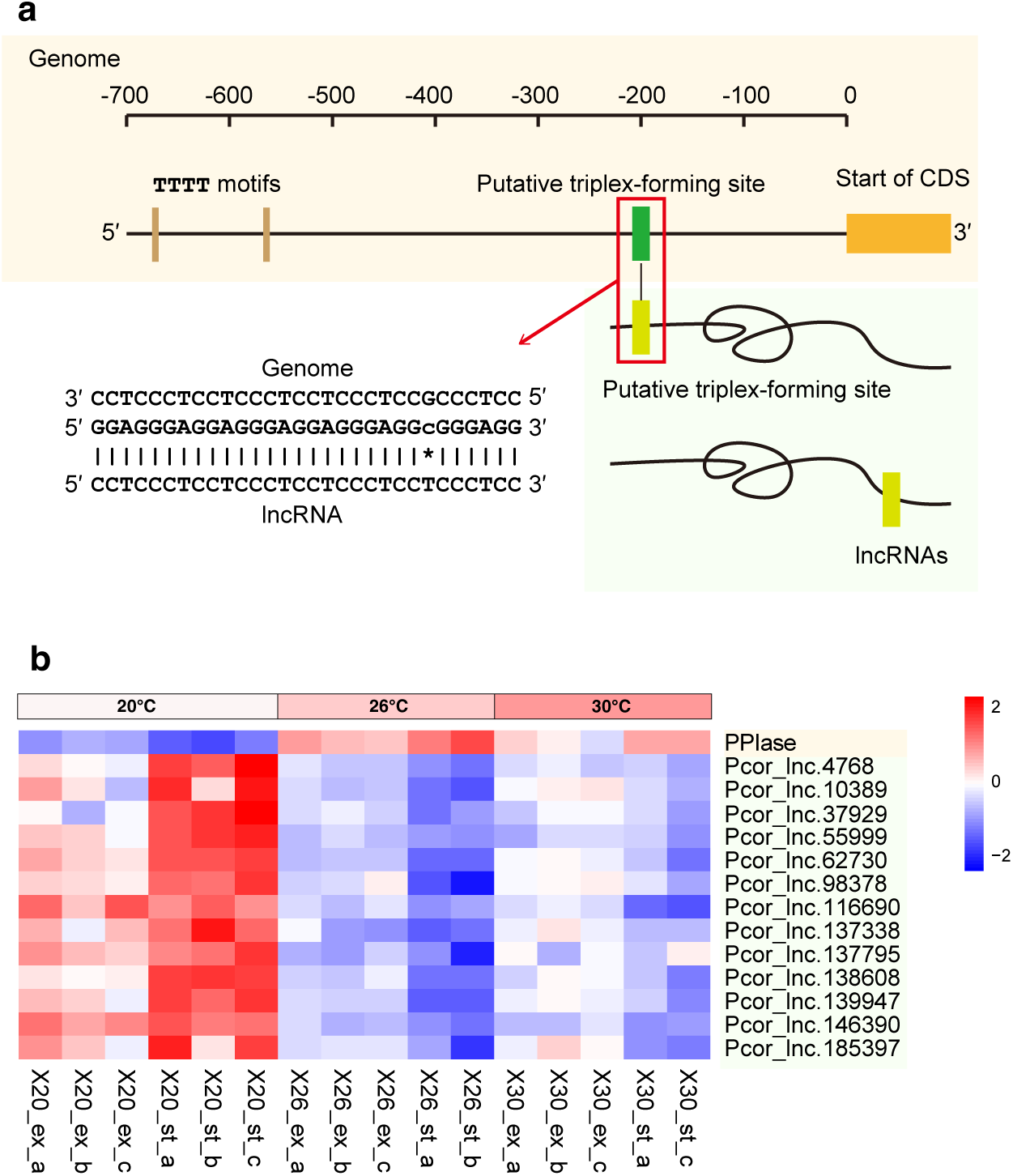
Hypothesised lncRNA regulation of the transcription of PPIase gene in *P. cordatum*, showing (a) the putative promoter region upstream of the protein-coding sequence and the predicted binding site for the formation of triple-helical structure, and (b) contrasting expression pattern of the up-regulated gene encoding PPIase and the 13 interacting lncRNAs that are down-regulated.

Notably, a TTTT motif was identified at 680 bp and 568 bp upstream of the coding sequence, which has been described as potential core promoter motif in dinoflagellates in lieu of the typical TATA-box in eukaryotes (Lin et al. 2015), and a known binding site for the general transcription factor TFIID involved in the formation of the transcription preinitiation complex (TPC) in dinoflagellates (Zaheri et al. 2022). TPC is a protein complex essential for transcription of protein-coding genes in eukaryotes (Greber and Nogales 2019); in human, triple-helical DNA structure at a promoter region is known to yield non-functional TPC, thus inhibiting transcription (Jain et al. 2010). We hypothesise that the expression of lncRNAs interacting with the putative promoter region of the PPIase-encoding gene in *P. cordatum* would similarly impede TPC formation, thus suppressing the transcription of the PPIase gene; likewise, the gene will be transcribed when the interacting lncRNAs were suppressed. Importantly, PPIase is known to form a heterocomplex with the 90-kDa heat-shock protein, thereby regulating various biological processes including stress responses (Subin et al. 2016; Zgajnar et al. 2019). This likely explains the up-regulation of this gene in *P. cordatum* during heat stress, supported by expression data across all sample replicates at 26°C and at 30°C relative to 20°C (Figure 5b), when the interacting lncRNAs were down-regulated.

### Concluding remarks

We present the first comprehensive survey of lncRNAs and their expression associated with the dinoflagellate stress response. Our results, based on high-quality transcriptome and genome data from dinoflagellates, reveal dinoflagellate lncRNAs are differentially expressed in response to stress, similarly to protein-coding genes. The conserved motifs for diverse lncRNAs in the genomes of three dinoflagellate taxa, combined with their co-expression with protein-coding genes, suggests a link between lncRNAs and gene expression and their functional role as regulatory elements for molecular responses in dinoflagellates. These results demonstrate the utility of *k*-mers in analysing complex, highly divergent genomic elements. We also discovered potential gene targets whose expression may be regulated by triplex-forming lncRNAs; these lncRNAs and their proposed functions are strong candidates for experimental validation. Knowledge of RNA-protein binding and RNA-DNA binding in dinoflagellates will further elucidate the possible functional domains (i.e. conserved structural features) in lncRNAs. Our results demonstrate how the inclusion of lncRNAs in transcriptome analyses provides novel insights into the molecular mechanisms that underpin RNA-based regulation of gene expression in dinoflagellates in the context of complex, atypical eukaryote genomes, that may extend more broadly to other microbial eukaryotes.

## Methods and Materials

### Data

We used published genome assemblies of *Cladocopium proliferum* SCF055 (Chen et al. 2022b), *Durusdinium trenchii* CCMP2556 (Dougan et al. 2022a) and *Prorocentrum cordatum* CCMP1329 (Dougan et al. 2022b), for which the associated transcriptome data are also available. For each of these taxa, a core transcriptome dataset, designed for heat stress experiments (Table 1), were carefully selected to minimise technical biases. For the C. proliferum dataset that consists of 24 samples (Camp et al. 2022), 16 representing T1 (early response) and TE (late response) were included (Table 1 and Table S10); the remaining eight were generated at control conditions that did not relate to heat stress.

### Identification of lncRNAs

We developed a customised workflow to identify putative lncRNAs using genome and transcriptome data of dinoflagellates. For each isolate, all raw RNA-Seq reads from the core transcriptome dataset were processed and adapter-trimmed using Fastp v0.23.2 (Chen et al. 2018), and assembled into transcripts *de novo* using Trinity v2.9.1 (Grabherr et al. 2011). For *C. proliferum*, due to the low data yield of the core dataset (Table 1), raw reads from other available transcriptome data (Levin et al. 2016; Chakravarti et al. 2020) (Table S10) were also included in this step to maximise recovery. The resulted transcripts were then mapped to their corresponding genome sequences with minimap v2.24 (Li 2018) for which the code was modified to recognise alternative splice sites of dinoflagellate genes, and further assembled using PASA v2.3.3 (Haas et al. 2003). Because the RNA-Seq data are generated from polyadenylated transcripts, these PASA-assembled transcripts represent polyadenylated protein-coding genes and lncRNAs. Protein-coding genes, i.e. transcripts that overlap with one or more exonic regions, were identified and removed. The coding potential of the remaining transcripts were assessed using three methods: CPC2 (Kang et al. 2017) at default setting, and FEELnc (Wucher et al. 2017) independently at *-m intergenic* and *-m shuffle* modes. Transcripts classified as non-coding by all three methods were retained. We then filtered out transcripts that share significant sequence similarity to known Pfam protein domains (BLASTx *p ≤*10^-5^); the remainder represent potential lncRNA candidates. To reduce redundancy, overlapping sequences were clustered based on genome location, among which the longest sequence was selected as the representative lncRNA from each cluster. To identify lncRNAs at high confidence for each isolate, we mapped the RNA-Seq reads from the core transcriptome dataset to the corresponding genome sequences using HISAT v2.2.1 (Kim et al. 2019), and counted the number of uniquely mapped reads using *featureCounts* (Liao et al. 2014). Only lncRNA candidates (identified above) supported by 10 or more samples (each with 10 or more uniquely mapped reads), were considered as the high-confident putative lncRNAs and used for subsequent analysis; 48,039 putative lncRNAs were identified this way from all three taxa (Table 2).

### Identification of homologous lncRNA based on overall sequence similarity

Among the 48,039 lncRNAs identified above, we identified homologous sets based on sequence similarity using OrthoFinder v2.3.8 (Emms and Kelly 2019). To identify putative functions of lncRNAs, we searched the 480,349 lncRNAs against the high-quality curated database of human lncRNAs, FANTOM CAT (89,998 sequences tagged as *lv3_robust*) (Hon et al. 2017) and all lncRNA sequences in the NCBI nr database (17,107; acquired on 15 August 2022), using BLASTn v2.11.0+ (Camacho et al. 2009).

### Identification of homologous lncRNA based on conserved motifs

We used seekr (Kirk et al. 2018) to group the 48,039 putative lncRNAs into ten clusters based on shared *k*-mers (at *k* = 4). Specifically, we used *seekr_k-mer_count* to count the number of distinct 4-mers in each lncRNA and normalised the count using *seekr_norm_vector* against 4-mers from all lncRNAs as background. The normalised 4-mer profiles were then used to calculate an adjacency matrix using *seekr_pearson* for each possible pair of lncRNAs. Based on hierarchical clustering, we used fcluster (*t=10, criterion=‘maxclust’*) available from the SciPy Python library (Virtanen et al. 2020) to group the lncRNAs into 10 clusters. This method is more scalable to the large datasets we used in this study compared to the Louvain algorithm used in Kirk et al. (2018), and was found to yield similar results in the same study.

### Analysis of differential expression of lncRNAs

For each core transcriptome dataset, filtered read count for transcripts (including both lncRNAs and genes) were normalised using the *getVarianceStabilizedData* implemented in DESeq2 (Love et al. 2014). To assess variation of samples within the dataset, the normalised counts were used to derive pairwise Euclidean distance between samples, from which the samples were grouped based on hierarchical clustering. Samples that did not group with other replicates were considered outliers and excluded from downstream analysis. We used DESeq2 (Love et al. 2014) to identify differentially expressed transcripts (i.e. lncRNAs and genes) related to heat stress, lifestyle, and/or growth phase. We consider transcripts with an adjusted (Benjamini and Hochberg 1995) *p* ≤ 0.01 and absolute (log_2_[fold-change]) ≥ 2 to be differentially expressed.

### Co-expression analysis

We performed weighted gene co-expression network analysis (WGCNA) (Langfelder and Horvath 2008) to assess transcript co-expression. The soft-thresholding power parameter (*T*), which assigns weighting of transcript co-expression, was carefully selected for each core transcriptome dataset. This was guided by the scale-free topology fit index (*I*), calculated from the normalised read counts of transcripts using the *pickSoftThreshold* function. We used *T* = 18 in all three datasets for different reasons. For *C. proliferum* and *P. cordatum*, *T* = 18 is the smallest value that gave rise to *I* > 0.8 (representing a good fit of the network). For *D. trenchii*, *I* was < 0.8 for *T* between 10 and 30; we therefore used *T* = 18 as recommended for signed networks with fewer than 20 samples (https://horvath.genetics.ucla.edu/html/CoexpressionNetwork/Rpackages/WGCNA/faq.html).

Next, we calculated the topological overlap matrix for signed network using *bicorr* correlation (*maxPOutliers=0.1, pearsonFallback=“individual”*) to assess co-expression similarity and adjacency, from which the dissimilarity was used to group the transcripts using hierarchical clustering (*method=complete*). Highly co-expressed transcripts were identified by “cutting” the branches using *dynamicTreeCut* (*deepSplit=2, pamRespectsDendro=FALSE, minModuleSize=30*). We then clustered module eigengenes (*method=average*) and merged those with high expression correlation (eigengene correlation ≥ 0.9).

To assess the correlation between k-mer clustering independently for each WGCNA module, we tested whether lncRNAs from any of the 10 k-mer clusters were significantly over– or under-represented using Fisher’s exact test (false discovery rate ≤ 0.01; two-tailed). To assess conserved motifs among a set of lncRNAs, we used MEME v5.5.3 (-dna –nmotifs 3) (Bailey et al. 2015).

### Identification of lncRNA targets

We used Triplexator (Buske et al. 2012) to search for possible formation of triple-helical structures between a lncRNA and the putative promoter region of a gene following the Hoogsteen base pairing rules. We defined putative promoter regions as 1Kb upstream of a coding sequence following Lin et al. (2015). Triplexator was run requiring a minimum length of 20bp for the binding region and no more than 5% mismatches, using the options *-l 20 –e 5 –fr on –mrl 7 –mrp 3 –dc 5 –of 1*.

## Author contributions

Conceptualisation, YC, KED, and CXC; methodology, YC, KED, QN, and CXC; formal analysis, YC; investigation, YC, KED; writing—original draft preparation, YC; writing— review and editing, YC, KED, QN, DB, and CXC; visualisation, YC; supervision, KED, QN, DB, CXC; funding acquisition, DB and CXC. All authors have read and agreed to the final version of the manuscript.

## Supporting information

Supplementary Tables S1 through S10

## Acknowledgements

This project is supported by high-performance computing facilities at the National Computational Infrastructure (NCI) National Facility systems through the NCI Merit Allocation Scheme (Project d85) awarded to CXC, the University of Queensland Research Computing Centre, and computing facility at the Australian Centre for Ecogenomics, School of Chemistry and Molecular Biosciences at the University of Queensland.

## References

1. Antonov IV, Mazurov E, Borodovsky M, Medvedeva YA (2019). Prediction of lncRNAs and their interactions with nucleic acids: benchmarking bioinformatics tools. Brief. Bioinform. 20:551–564.

2. Aranda M, Li Y, Liew YJ, Baumgarten S, Simakov O, Wilson MC, Piel J, Ashoor H, Bougouffa S, Bajic VB (2016). Genomes of coral dinoflagellate symbionts highlight evolutionary adaptations conducive to a symbiotic lifestyle. Sci. Rep. 6:39734.

3. Asai DJ, Wilkes DE (2004). The dynein heavy chain family. J. Eukaryot. Microbiol. 51:23–29.

4. Babu MM, Luscombe NM, Aravind L, Gerstein M, Teichmann SA (2004). Structure and evolution of transcriptional regulatory networks. Curr. Opin. Struct. Biol. 14:283–291.

5. Bailey TL, Johnson J, Grant CE, Noble WS (2015). The MEME Suite. Nucleic Acids Res. 43:W39–W49.

6. Barshis DJ, Ladner JT, Oliver TA, Palumbi SR (2014). Lineage-specific transcriptional profiles of *Symbiodinium* spp. unaltered by heat stress in a coral host. Mol. Biol. Evol. 31:1343–1352.

7. Bartel DP (2009). MicroRNAs: target recognition and regulatory functions. Cell 136:215–233.

8. Baumgarten S, Bayer T, Aranda M, Liew YJ, Carr A, Micklem G, Voolstra CR (2013). Integrating microRNA and mRNA expression profiling in *Symbiodinium microadriaticum*, a dinoflagellate symbiont of reef-building corals. BMC Genomics 14:704.

9. Bayer T, Aranda M, Sunagawa S, Yum LK, DeSalvo MK, Lindquist E, Coffroth MA, Voolstra CR, Medina M (2012). *Symbiodinium* transcriptomes: genome insights into the dinoflagellate symbionts of reef-building corals. PLoS ONE 7:e35269.

10. Beauchemin M, Roy S, Daoust P, Dagenais-Bellefeuille S, Bertomeu T, Letourneau L, Lang BF, Morse D (2012). Dinoflagellate tandem array gene transcripts are highly conserved and not polycistronic. Proc. Natl. Acad. Sci. U. S. A. 109:15793–15798.

11. Bellantuono AJ, Dougan KE, Granados-Cifuentes C, Rodriguez-Lanetty M (2019). Free-living and symbiotic lifestyles of a thermotolerant coral endosymbiont display profoundly distinct transcriptomes under both stable and heat stress conditions. Mol. Ecol. 28:5265–5281.

12. Benjamini Y, Hochberg Y (1995). Controlling the false discovery rate: a practical and powerful approach to multiple testing. J. R. Stat. Sco. Ser. B 57:289–300.

13. Buske FA, Bauer DC, Mattick JS, Bailey TL (2012). Triplexator: detecting nucleic acid triple helices in genomic and transcriptomic data. Genome Res. 22:1372–1381.

14. Butler CC, Turnham KE, Lewis AM, Nitschke MR, Warner ME, Kemp DW, Hoegh-Guldberg O, Fitt WK, van Oppen MJH, LaJeunesse TC (2023). Formal recognition of host-generalist species of dinoflagellate (*Cladocopium*, Symbiodiniaceae) mutualistic with Indo-Pacific reef corals. J. Phycol.:doi:10.1111/jpy.13340.

15. Camacho C, Coulouris G, Avagyan V, Ma N, Papadopoulos J, Bealer K, Madden TL (2009). BLAST+: architecture and applications. BMC Bioinformatics 10:421.

16. Camp EF, Kahlke T, Signal B, Oakley CA, Lutz A, Davy SK, Suggett DJ, Leggat WP (2022). Proteome metabolome and transcriptome data for three Symbiodiniaceae under ambient and heat stress conditions. Sci. Data 9:153.

17. Cerase A, Pintacuda G, Tattermusch A, Avner P (2015). Xist localization and function: new insights from multiple levels. Genome Biol. 16:166.

18. Chakravarti LJ, Buerger P, Levin RA, van Oppen MJH (2020). Gene regulation underpinning increased thermal tolerance in a laboratory-evolved coral photosymbiont. Mol. Ecol. 29:1684–1703.

19. Chen C-K, Yu C-P, Li S-C, Wu S-M, Lu M-YJ, Chen Y-H, Chen D-R, Ng CS, Ting C-T, Li W-H (2017). Identification and evolutionary analysis of long non-coding RNAs in zebra finch. BMC Genomics 18:117.

20. Chen S, Zhou Y, Chen Y, Gu J (2018). fastp: an ultra-fast all-in-one FASTQ preprocessor. Bioinformatics 34:i884–i890.

21. Chen T, Liu Y, Song S, Bai J, Li C (2022a). Full-length transcriptome analysis of the bloom-forming dinoflagellate *Akashiwo sanguinea* by single-molecule real-time sequencing. Front. Microbiol. 13:993914.

22. Chen Y, González-Pech RA, Stephens TG, Bhattacharya D, Chan CX (2020). Evidence that inconsistent gene prediction can mislead analysis of dinoflagellate genomes. J. Phycol. 56:6–10.

23. Chen Y, Shah S, Dougan KE, van Oppen MJ, Bhattacharya D, Chan CX (2022b). Improved *Cladocopium goreaui* genome assembly reveals features of a facultative coral symbiont and the complex evolutionary history of dinoflagellate genes. Microorganisms 10:1662.

24. Diederichs S (2014). The four dimensions of noncoding RNA conservation. Trends Genet. 30:121–123.

25. Dougan KE, Bellantuono AJ, Kahlke T, Abbriano RM, Chen Y, Shah S, Granados-Cifuentes C, van Oppen MJ, Bhattacharya D, Suggett DJ (2022a). Whole-genome duplication in an algal symbiont serendipitously confers thermal tolerance to corals. bioRxiv:2022.2004.2010.487810.

26. Dougan KE, Deng Z-L, Woehlbrand L, Reuse C, Bunk B, Chen Y, Hartlich J, Hiller K, John U, Kalvelage J (2022b). Multi-omics analysis reveals the molecular response to heat stress in a “red tide” dinoflagellate. bioRxiv:2022.2007.2025.501386.

27. Emms DM, Kelly S (2019). OrthoFinder: phylogenetic orthology inference for comparative genomics. Genome Biol. 20:238.

28. Fan Y, Li J, Yang Q, Gong C, Gao H, Mao Z, Yuan X, Zhu S, Xue Z (2019). Dysregulated long non-coding RNAs in Parkinson’s disease contribute to the apoptosis of human neuroblastoma cells. Front. Neurosci. 13:1320.

29. González-Pech RA, Stephens TG, Chen Y, Mohamed AR, Cheng Y, Shah S, Dougan KE, Fortuin MDA, Lagorce R, Burt DW, et al. (2021). Comparison of 15 dinoflagellate genomes reveals extensive sequence and structural divergence in family Symbiodiniaceae and genus *Symbiodinium*. BMC Biol. 19:73.

30. Grabherr MG, Haas BJ, Yassour M, Levin JZ, Thompson DA, Amit I, Adiconis X, Fan L, Raychowdhury R, Zeng Q (2011). Trinity: reconstructing a full-length transcriptome without a genome from RNA-Seq data. Nat. Biotechnol. 29:644.

31. Greber BJ, Nogales E (2019). The structures of eukaryotic transcription pre-initiation complexes and their functional implications. Subcell. Biochem. 93:143–192.

32. Grote P, Herrmann BG (2013). The long non-coding RNA Fendrr links epigenetic control mechanisms to gene regulatory networks in mammalian embryogenesis. RNA Biol. 10:1579–1585.

33. Grote P, Wittler L, Hendrix D, Koch F, Währisch S, Beisaw A, Macura K, Bläss G, Kellis M, Werber M (2013). The tissue-specific lncRNA Fendrr is an essential regulator of heart and body wall development in the mouse. Dev. Cell 24:206–214.

34. Haas BJ, Delcher AL, Mount SM, Wortman JR, Smith Jr RK, Hannick LI, Maiti R, Ronning CM, Rusch DB, Town CD (2003). Improving the *Arabidopsis* genome annotation using maximal transcript alignment assemblies. Nucleic Acids Res. 31:5654–5666.

35. Hombach S, Kretz M (2016). Non-coding RNAs: classification, biology and functioning. Adv. Exp. Med. Biol. 937:3–17.

36. Hon C-C, Ramilowski JA, Harshbarger J, Bertin N, Rackham OJL, Gough J, Denisenko E, Schmeier S, Poulsen TM, Severin J, et al. (2017). An atlas of human long non-coding RNAs with accurate 5’ ends. Nature 543:199–204.

37. Hou Y, Lin S (2009). Distinct gene number-genome size relationships for eukaryotes and non-eukaryotes: gene content estimation for dinoflagellate genomes. PLoS ONE 4:e6978.

38. Ip JY, Nakagawa S (2012). Long non-coding RNAs in nuclear bodies. Dev. Growth Differ. 54:44–54.

39. Jain A, Magistri M, Napoli S, Carbone GM, Catapano CV (2010). Mechanisms of triplex DNA-mediated inhibition of transcription initiation in cells. Biochimie 92:317–320.

40. John U, Lu Y, Wohlrab S, Groth M, Janouškovec J, Kohli GS, Mark FC, Bickmeyer U, Farhat S, Felder M, et al. (2019). An aerobic eukaryotic parasite with functional mitochondria that likely lacks a mitochondrial genome. Sci. Adv. 5:eaav1110.

41. Johnsson P, Lipovich L, Grandér D, Morris KV (2014). Evolutionary conservation of long non-coding RNAs; sequence, structure, function. Biochim. Biophys. Acta Biochim Biophys Acta 1840:1063–1071.

42. Kang HC, Jeong HJ, Park SA, Ok JH, You JH, Eom SH, Park EC, Jang SH, Lee SY (2021). Comparative transcriptome analysis of the phototrophic dinoflagellate *Biecheleriopsis adriatica* grown under optimal temperature and cold and heat stress. Front. Mar. Sci. 8:761095.

43. Kang Y-J, Yang D-C, Kong L, Hou M, Meng Y-Q, Wei L, Gao G (2017). CPC2: a fast and accurate coding potential calculator based on sequence intrinsic features. Nucleic Acids Res. 45:W12–W16.

44. Kaur P, Liu F, Tan JR, Lim KY, Sepramaniam S, Karolina DS, Armugam A, Jeyaseelan K (2013). Non-coding RNAs as potential neuroprotectants against ischemic brain injury. Brain Sci. 3:360–395.

45. Kim D, Paggi JM, Park C, Bennett C, Salzberg SL (2019). Graph-based genome alignment and genotyping with HISAT2 and HISAT-genotype. Nat. Biotechnol. 37:907–915.

46. Kirk JM, Kim SO, Inoue K, Smola MJ, Lee DM, Schertzer MD, Wooten JS, Baker AR, Sprague D, Collins DW, et al. (2018). Functional classification of long non-coding RNAs by k-mer content. Nat. Genet. 50:1474–1482.

47. Kopp F, Mendell JT (2018). Functional classification and experimental dissection of long noncoding RNAs. Cell 172:393–407.

48. Langfelder P, Horvath S (2008). WGCNA: an R package for weighted correlation network analysis. BMC Bioinformatics 9:559.

49. Leisegang MS, Bains JK, Seredinski S, Oo JA, Krause NM, Kuo C-C, Günther S, Sentürk Cetin N, Warwick T, Cao C, et al. (2022). HIF1α-AS1 is a DNA:DNA:RNA triplex-forming lncRNA interacting with the HUSH complex. Nat. Commun. 13:6563.

50. Levin RA, Beltran VH, Hill R, Kjelleberg S, McDougald D, Steinberg PD, van Oppen MJH (2016). Sex, scavengers, and chaperones: transcriptome secrets of divergent *Symbiodinium* thermal tolerances. Mol. Biol. Evol. 33:2201–2215.

51. Li D, Yang MQ (2017). Identification and characterization of conserved lncRNAs in human and rat brain. BMC Bioinformatics 18:31–38.

52. Li H (2018). Minimap2: pairwise alignment for nucleotide sequences. Bioinformatics 34:3094–3100.

53. Li Y, Syed J, Sugiyama H (2016). RNA-DNA triplex formation by long noncoding RNAs. Cell Chem. Biol. 23:1325–1333.

54. Liao Y, Smyth GK, Shi W (2014). featureCounts: an efficient general purpose program for assigning sequence reads to genomic features. Bioinformatics 30:923–930.

55. Liew YJ, Li Y, Baumgarten S, Voolstra CR, Aranda M (2017). Condition-specific RNA editing in the coral symbiont *Symbiodinium microadriaticum*. PLoS Genet. 13:e1006619.

56. Lin S (2011). Genomic understanding of dinoflagellates. Res. Microbiol. 162:551–569.

57. Lin S, Cheng S, Song B, Zhong X, Lin X, Li W, Li L, Zhang Y, Zhang H, Ji Z (2015). The *Symbiodinium kawagutii* genome illuminates dinoflagellate gene expression and coral symbiosis. Science 350:691–694.

58. Lizio M, Harshbarger J, Shimoji H, Severin J, Kasukawa T, Sahin S, Abugessaisa I, Fukuda S, Hori F, Ishikawa-Kato S, et al. (2015). Gateways to the FANTOM5 promoter level mammalian expression atlas. Genome Biol. 16:22.

59. Lo R, Dougan KE, Chen Y, Shah S, Bhattacharya D, Chan CX (2022). Alignment-free analysis of whole-genome sequences from Symbiodiniaceae reveals different phylogenetic signals in distinct regions. Front. Plant Sci. 13:815714.

60. Love MI, Huber W, Anders S (2014). Moderated estimation of fold change and dispersion for RNA-seq data with DESeq2. Genome Biol. 15:550.

61. Lowe CD, Mello LV, Samatar N, Martin LE, Montagnes DJS, Watts PC (2011). The transcriptome of the novel dinoflagellate *Oxyrrhis marina* (Alveolata: Dinophyceae): response to salinity examined by 454 sequencing. BMC Genomics 12:519.

62. Martianov I, Ramadass A, Serra Barros A, Chow N, Akoulitchev A (2007). Repression of the human dihydrofolate reductase gene by a non-coding interfering transcript. Nature 445:666–670.

63. Mattick JS, Amaral PP, Carninci P, Carpenter S, Chang HY, Chen LL, Chen R, Dean C, Dinger ME, Fitzgerald KA, et al. (2023). Long non-coding RNAs: definitions, functions, challenges and recommendations. Nat. Rev. Mol. Cell Biol. 24:430–447.

64. Moison M, Pacheco JM, Lucero L, Fonouni-Farde C, Rodríguez-Melo J, Mansilla N, Christ A, Bazin J, Benhamed M, Ibañez F, et al. (2021). The lncRNA APOLO interacts with the transcription factor WRKY42 to trigger root hair cell expansion in response to cold. Mol. Plant 14:937–948.

65. Moustafa A, Evans AN, Kulis DM, Hackett JD, Erdner DL, Anderson DM, Bhattacharya D (2010). Transcriptome profiling of a toxic dinoflagellate reveals a gene-rich protist and a potential impact on gene expression due to bacterial presence. PLoS ONE 5:e9688.

66. Palos K, Nelson Dittrich AC, Yu L, Brock JR, Railey CE, Wu H-YL, Sokolowska E, Skirycz A, Hsu PY, Gregory BD, et al. (2022). Identification and functional annotation of long intergenic non-coding RNAs in Brassicaceae. Plant Cell 34:3233–3260.

67. Roy S, Beauchemin M, Dagenais-Bellefeuille S, Letourneau L, Cappadocia M, Morse D (2014). The *Lingulodinium* circadian system lacks rhythmic changes in transcript abundance. BMC Biol. 12:107.

68. Sentürk Cetin N, Kuo C-C, Ribarska T, Li R, Costa IG, Grummt I (2019). Isolation and genome-wide characterization of cellular DNA:RNA triplex structures. Nucleic Acids Res. 47:2306–2321.

69. Shi X, Lin X, Li L, Li M, Palenik B, Lin S (2017). Transcriptomic and microRNAomic profiling reveals multi-faceted mechanisms to cope with phosphate stress in a dinoflagellate. ISME J. 11:2209–2218.

70. Shoguchi E, Beedessee G, Tada I, Hisata K, Kawashima T, Takeuchi T, Arakaki N, Fujie M, Koyanagi R, Roy MC (2018). Two divergent *Symbiodinium* genomes reveal conservation of a gene cluster for sunscreen biosynthesis and recently lost genes. BMC Genomics 19:458.

71. Shoguchi E, Shinzato C, Kawashima T, Gyoja F, Mungpakdee S, Koyanagi R, Takeuchi T, Hisata K, Tanaka M, Fujiwara M (2013). Draft assembly of the *Symbiodinium minutum* nuclear genome reveals dinoflagellate gene structure. Curr. Biol. 23:1399–1408.

72. Stephens TG, González-Pech RA, Cheng Y, Mohamed AR, Burt DW, Bhattacharya D, Ragan MA, Chan CX (2020). Genomes of the dinoflagellate *Polarella glacialis* encode tandemly repeated single-exon genes with adaptive functions. BMC Biol. 18:56.

73. Subin CS, Pradeep MA, Vijayan KK (2016). FKBP-type peptidyl-prolyl cis-trans isomerase from thermophilic microalga, Scenedesmus sp.: molecular characterisation and demonstration of acquired salinity and thermotolerance in E. coli by recombinant expression. J. Appl. Phycol. 28:3307–3315.

74. Vallon O (2005). Chlamydomonas immunophilins and parvulins: survey and critical assessment of gene models. Eukaryot. Cell 4:230–241.

75. Virtanen P, Gommers R, Oliphant TE, Haberland M, Reddy T, Cournapeau D, Burovski E, Peterson P, Weckesser W, Bright J, et al. (2020). SciPy 1.0: fundamental algorithms for scientific computing in Python. Nat. Methods 17:261–272.

76. Wang X, Niu X, Chen Y, Sun Z, Han A, Lou X, Ge J, Li X, Yang Y, Jian J, et al. (2019). Transcriptome sequencing of a toxic dinoflagellate, *Karenia mikimotoi* subjected to stress from solar ultraviolet radiation. Harmful Algae 88:101640.

77. Wisecaver JH, Hackett JD (2011). Dinoflagellate genome evolution. Annu. Rev. Microbiol. 65:369–387.

78. Wucher V, Legeai F, Hédan B, Rizk G, Lagoutte L, Leeb T, Jagannathan V, Cadieu E, David A, Lohi H, et al. (2017). FEELnc: a tool for long non-coding RNA annotation and its application to the dog transcriptome. Nucleic Acids Res. 45:e57.

79. Yu L, Zhang Y, Li M, Wang C, Lin X, Li L, Shi X, Guo C, Lin S (2020). Comparative metatranscriptomic profiling and microRNA sequencing to reveal active metabolic pathways associated with a dinoflagellate bloom. Sci. Total Environ. 699:134323.

80. Zaheri B, Morse D (2021). Assessing nucleic acid binding activity of four dinoflagellate cold shock domain proteins from *Symbiodinium kawagutii* and *Lingulodinium polyedra*. BMC Mol. Cell Biol. 22:27.

81. Zaheri B, Veilleux-Trinh C, Morse D (2022). A dinoflagellate TBP-like factor activates transcription from a TTTT-box in yeast. J. Phycol. 58:343–346.

82. Zgajnar NR, De Leo SA, Lotufo CM, Erlejman AG, Piwien-Pilipuk G, Galigniana MD (2019). Biological Actions of the Hsp90-binding Immunophilins FKBP51 and FKBP52. Biomolecules 9:52.

83. Zhang H, Hou Y, Miranda L, Campbell DA, Sturm NR, Gaasterland T, Lin S (2007). Spliced leader RNA trans-splicing in dinoflagellates. Proc. Natl. Acad. Sci. U. S. A. 104:4618–4623.

