## Supplementary Figures S1 and S2 for "Genome-wide transcriptome analysis reveals the diversity and function of long non-coding RNAs in dinoflagellates"

**a**

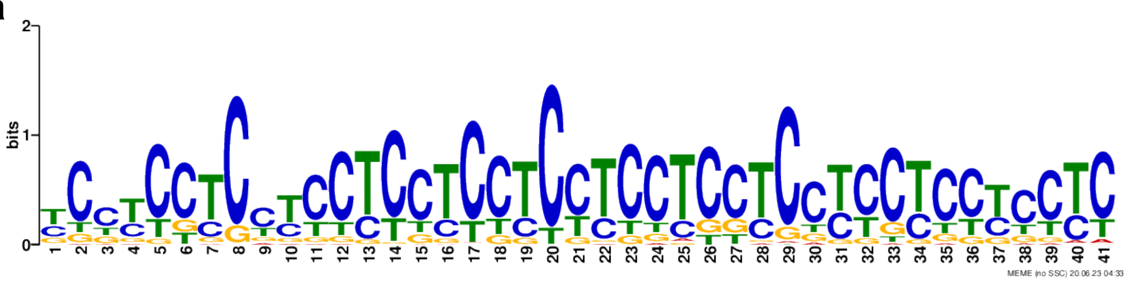


The most frequent motif in *k*-mer clusters 5 and 6, in M7 and M8

```
MOTIF YCCYCCTCCYCCTCCTCCTCCTCCTCCTCCYCCYCCYCCTC
width = 41  sites = 299  llr = 5793  E-value = 2.1e-162
```

**b**

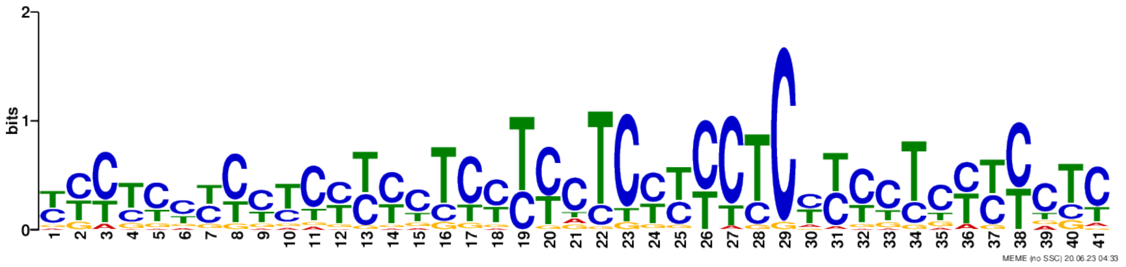


The most frequent motif in *k*-mer clusters 2, 3, and 10, in M46 and M49

```
MOTIF YYCYCYYCYYCYYYCTCYYYCTCYYYCTCYYYCTCYYYCTC
width = 41  sites = 996  llr = 15952  E-value = 2.9e-082
```

**Figure S1.** The conserved CT-rich motifs lncRNAs in *k*-mer clusters that were found to be significantly overrepresented in distinct WGCNA modules.

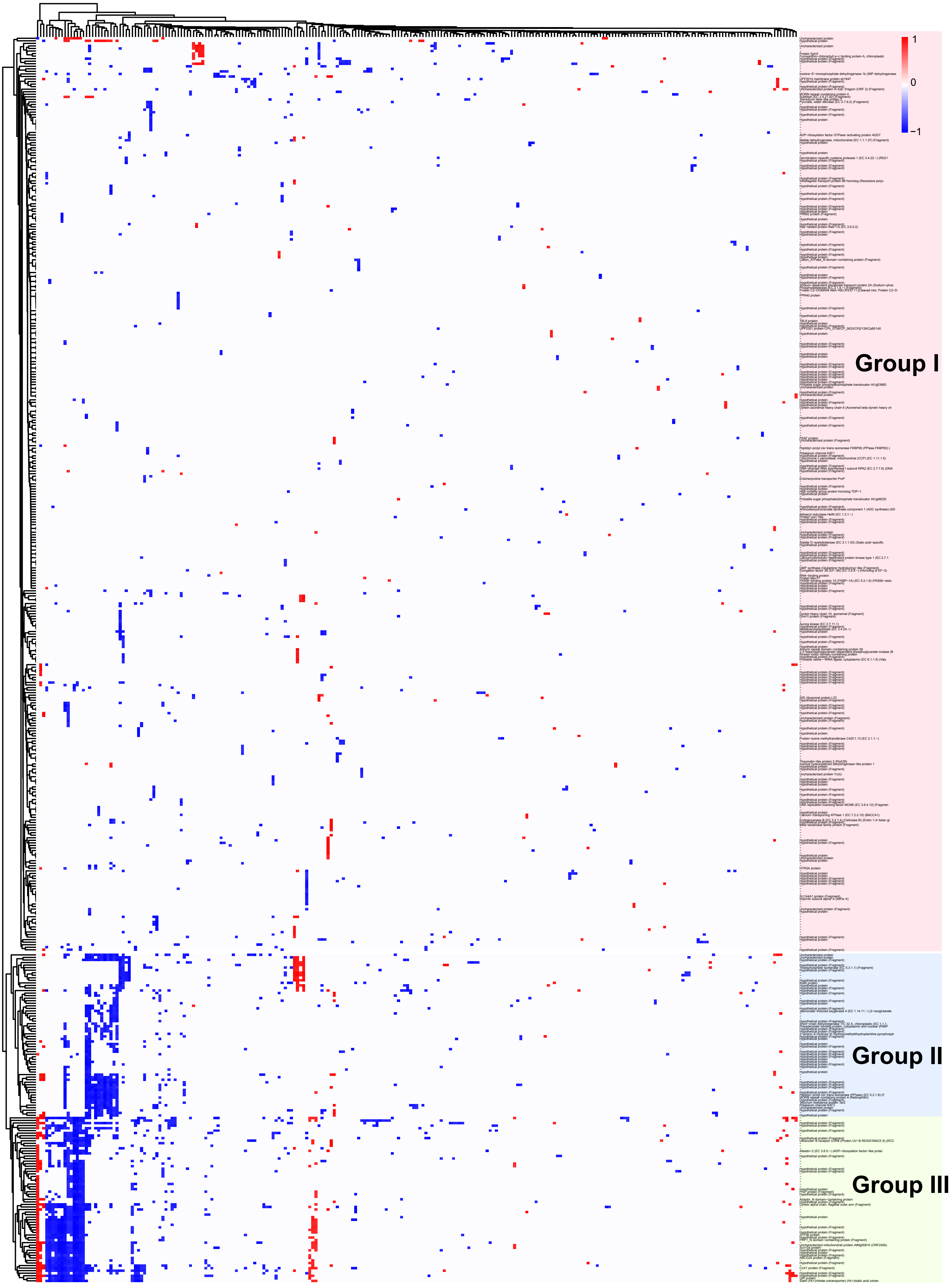


**Figure S2.** Correlation of expression of differentially expressed *P. cordatum* genes under heat stress (30 versus 20°C) with the expression of their interacting lncRNAs.
